# Engineering an Enzymatically Active Granular Matrix for On-Chip Modeling of Bone-Like Mineralization

**DOI:** 10.64898/2026.07.12.737035

**Authors:** Farhad Sanaei, Dina Zandieh, Daniel Hofman, Lucas Joziasse, Jeroen JJP van den Beucken, Sander CG Leeuwenburgh, Mani Diba

**Author notes:** Institute for Complex Molecular Systems, Eindhoven University of Technology, Eindhoven 5600 MB, The Netherlands.

## Abstract

Controlled biomineralization is central to engineering physiologically relevant hard-tissue models, yet achieving spatially organized, three-dimensional (3D) mineral deposition in microfluidic on-chip systems remains challenging. While cell-based bone-on-chip platforms offer biological complexity, they intrinsically couple mineral initiation to confounding factors such as matrix remodeling and paracrine signaling, obscuring the earliest biochemical drivers of nucleation. Drawing inspiration from bottom-up synthetic biology, we engineered an enzymatically active granular matrix that recapitulates a key osteogenic function within a perfusable 3D microenvironment. Alkaline phosphatase (ALP), the key driver of native bone formation, was covalently conjugated to poly(ethylene glycol)-based (PEG) microgels via thiol–ene photochemistry, retaining over 90% enzymatic activity after 48 h. These monodisperse microgels were assembled into a jammed, perfusable matrix within an on-chip chamber, enabling independent control over enzyme loading and substrate delivery. The system supported rapid in situ mineralization (24–48 h), yielding a carbonated, calcium-deficient, apatite-like phase characteristic of early-stage bone mineral. We demonstrate that the spatial 3D localization of enzymatic activity to discrete microscale compartments, coupled with interstitial perfusion, enables localized and near-physiological mineral formation. This mechanistically defined, acellular platform provides a programmable foundation for investigating ALP-driven 3D mineralization and establishes a modular route toward hybrid biosynthetic models of (patho)physiological tissue mineralization.

## 1. Introduction

Bone mineralization is a tightly regulated process orchestrated by osteoblasts through spatially compartmentalized enzymatic activity.^[1,2]^ During bone formation, osteoblasts express alkaline phosphatase (ALP), an ectoenzyme localized to the cell membrane and matrix vesicles. ALP hydrolyzes organic phosphate esters to elevate local inorganic phosphate concentrations and promote calcium phosphate nucleation.^[3,4]^ Spatial localization is key for this process, as osteoblasts localize mineral initiation to defined sites rather than permitting nonspecific bulk precipitation by restricting ALP activity to membrane- and vesicle-confined microdomains.^[1]^ Mineral deposition is subsequently shaped by additional regulatory mechanisms, including matrix vesicle-mediated initiation, cell-controlled modulation of the ionic microenvironment, and extracellular matrix (ECM) templating by collagen and non-collagenous proteins.^[5]^ Disruption of this spatially controlled process contributes to pathological mineralization in disorders such as osteoporosis, osteomalacia, and hypophosphatasia.^[4,6,7]^ Mechanistic investigation of early-stage bone-like mineralization and the development of targeted therapies would therefore require in vitro platforms that faithfully recapitulate this spatially localized, ALP-driven process.

Recently, organ-on-chip systems have emerged as promising microphysiological systems that can recapitulate (patho)physiological processes under controlled conditions.^[8]^ More specifically, bone-on-chip systems have emerged as in vitro platforms for studying osteogenesis, matrix deposition, and mineralization. Accordingly, several cell-based bone-on-chip models have been developed recapitulating bone biology mainly within 2D and, more recently, 3D microenvironments.^[9]^ These systems offer important advantages, including microscale control over cellular complexity, tissue geometry, and culture conditions.^[9]^ However, their focus on recapitulating biological complexity also limits their utility for isolating the earliest biochemical drivers of mineral initiation. In particular, cell-based systems intrinsically couple enzymatically driven mineralization to confounding factors such as matrix remodeling, paracrine signaling, differentiation state, and donor-dependent variability, while also requiring prolonged culture periods that constrain experimentation throughput.^[9]^ As a result, it remains challenging to resolve the specific contribution of localized ALP activity to mineral nucleation and early phase bone-like mineralization in these platforms.

Acellular mineralization systems offer a complementary strategy to cell-based models by reducing biological variability and enabling greater control over physicochemical conditions.^[10]^ Direct incorporation of inorganic components such as calcium phosphates or bioactive glasses in in vitro models can reproduce mineral content, yet offers limited control over nucleation kinetics or spatial architecture of the mineral phase.^[10,11]^ ALP-mediated systems are particularly attractive as they reconstitute a core biochemical driver of bone mineralization under near-physiological conditions.^[12,13]^ Recent studies have incorporated enzyme-induced mineralization into i) hydrogels,^[12,14]^ ii) 3D-printed scaffolds^[15,16]^ and iii) onto implants^[17]^ to promote in situ mineral formation and enhance the mineralization capacity of engineered constructs. However, these efforts have primarily focused on material design rather than the recapitulation of spatially localized ALP activity in a controllable microenvironment. Consequently, current approaches do not yet enable compartmentalized ALP activity in a perfusable 3D environment.

Recent advances in bottom-up synthetic biology and cytomimetic materials highlight compartmentalization as a promising avenue for addressing this challenge.^[18]^ Protocells and prototissues have shown that biochemical reactions can be spatially localized to programmable microscale domains, enabling selective reconstruction of specific biological functions, while reducing confounding biological complexity.^[19–22]^ Cytomimetic mineralizing materials further suggest that localized catalytic environments and matrix properties jointly regulate calcium phosphate nucleation and further maturation of the mineral phase, achieving self-regulated calcification and decalcification within integrated protocell/matrix composites.^[18]^ However, such systems operate as static bulk constructs and are not designed to serve as modular, perfusable on-chip platforms for mechanistic modeling of ALP-driven mineral formation.

In recent years, granular hydrogels assembled from microgel building blocks have emerged as modular biomaterials for creating 3D cell culture matrices.^[23]^ Their inherent interparticle microporosity supports cell infiltration and molecular transport without requiring matrix degradation.^[24,25]^ Moreover, these materials can exhibit shear-thinning and self-healing behavior, enabling their injection or loading into confined geometries such as narrow channels.^[26]^ Together, these properties render granular hydrogels attractive 3D matrices for organ-on-chip systems. However, their capacity as engineered matrices for directing mineral formation in on-chip systems remains unexplored.

Here, we report an enzymatically active granular matrix for on-chip modeling of bone-like mineralization. To this end, we designed PEG-based microgels bearing covalently conjugated ALP enzymes as modular building blocks for 3D matrix assembly (**Figure 1a-b**). These microgels were generated by means of droplet microfluidics and formed via photoinitiated thiol-ene chemistry, enabling simultaneous hydrogel network formation and enzyme conjugation within the hydrogel phase. Upon physical jamming in a 3D microfluidic chamber, these building blocks assembled into a perfusable granular matrix in which ALP activity was spatially localized to discrete microscale domains (Figure 1c). This on-chip platform enables localized ALP-driven mineralization within a tightly controlled 3D environment (Figure 1d). Our results demonstrate that this system supports near-physiological, bone-like mineralization within 24-48 h, yields an apatite-like phase characteristic of early-stage bone mineral, and allows modulation of mineralization by variations in microgel design and in vitro incubation conditions. Taken together, this work establishes a mechanistically defined acellular platform for in vitro modeling of ALP-driven mineralization and provides a tunable foundation for future biosynthetic models of tissue mineralization.

**Figure 1.**
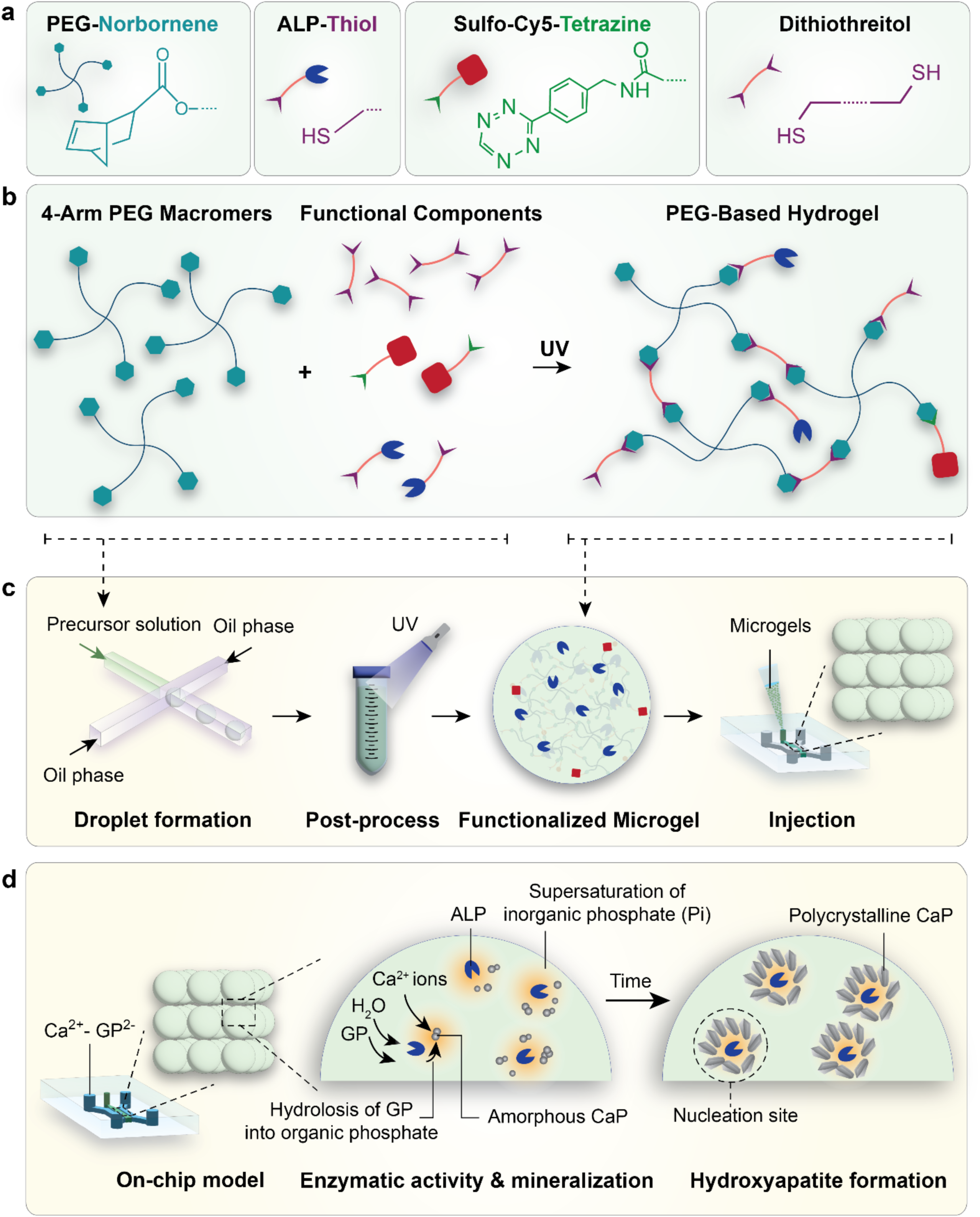
Experimental design of the enzymatically active granular matrix and 3D on-chip mineralization model. (a) Schematic of granular matrix building blocks and corresponding molecular structures. (b) Workflow for UV-mediated formation of hydrogels as an enzymatically active matrix. 4-arm PEG-norbornene (PEG-Nb) macromers are combined with dithiothreitol (DTT) as the crosslinker, thiolated alkaline phosphatase (SH-ALP) as the bioactive addictive, and Sulfo-Cyanine5-Tetrazine (Sulfo-Cy5-Tetrazine) as the fluorescent additive. Upon UV-exposure, photo-induced thiol-ene reaction yields a PEG-based hydrogel network with conjugated ALP. (c) Workflow for microgel fabrication and their on-chip 3D assembly. Microgels are generated via flow-focusing microfluidics by pinching the hydrogel precursor solution into microdroplets in an oil phase, followed by collection and UV curing. The formed microgels are then physically jammed into the microfluidic chip by injection into the central channel. (d) Schematic illustration of the resulting enzymatically active microgels enabling on-chip recapitulation of ALP-driven mineralization. ALP-mediated hydrolysis of glycerophosphate (GP) creates a local supersaturation of inorganic phosphate (Pi), triggering the deposition of amorphous calcium phosphate (ACP) in the presence of Ca^2+^ ions. Subsequent incubation of ACP at 37 °C leads to the maturation of these minerals into biomimetic crystalline hydroxyapatite.

## 2. Results and Discussion

### 2.1. Thiolation of ALP yields tunable thiol density with preserved enzymatic activity

Stable, long-term retention of enzymatic activity within the microgels is essential for sustained on-chip mineral formation, which requires covalent immobilization of ALP within the microgel polymer network to prevent leaching during perfusion. To this end, we employed photo-initiated thiol-ene click chemistry, as it offers a more rapid and selective reaction for hydrogel network cross-linking and ALP conjugation, compared to conventional protein conjugation strategies such as carbodiimide/NHS coupling.^[27,28]^ Successful conjugation of ALP to the microgels through thiol-ene reaction requires accessible thiol groups on the enzyme surface.^[29]^ Since bovine intestinal ALP employed herein possesses limited accessible free thiols for direct coupling,^[30]^ ALP was first thiolated through amine-reactive modification prior to immobilization, using Traut’s reagent (2-iminothiolane). To identify functionalization conditions that maximize functional thiol incorporation while preserving enzymatic activity, the effects of Traut’s reagent concentration, pH, and reaction time were systematically evaluated (**Figure 2**).

**Figure 2.**
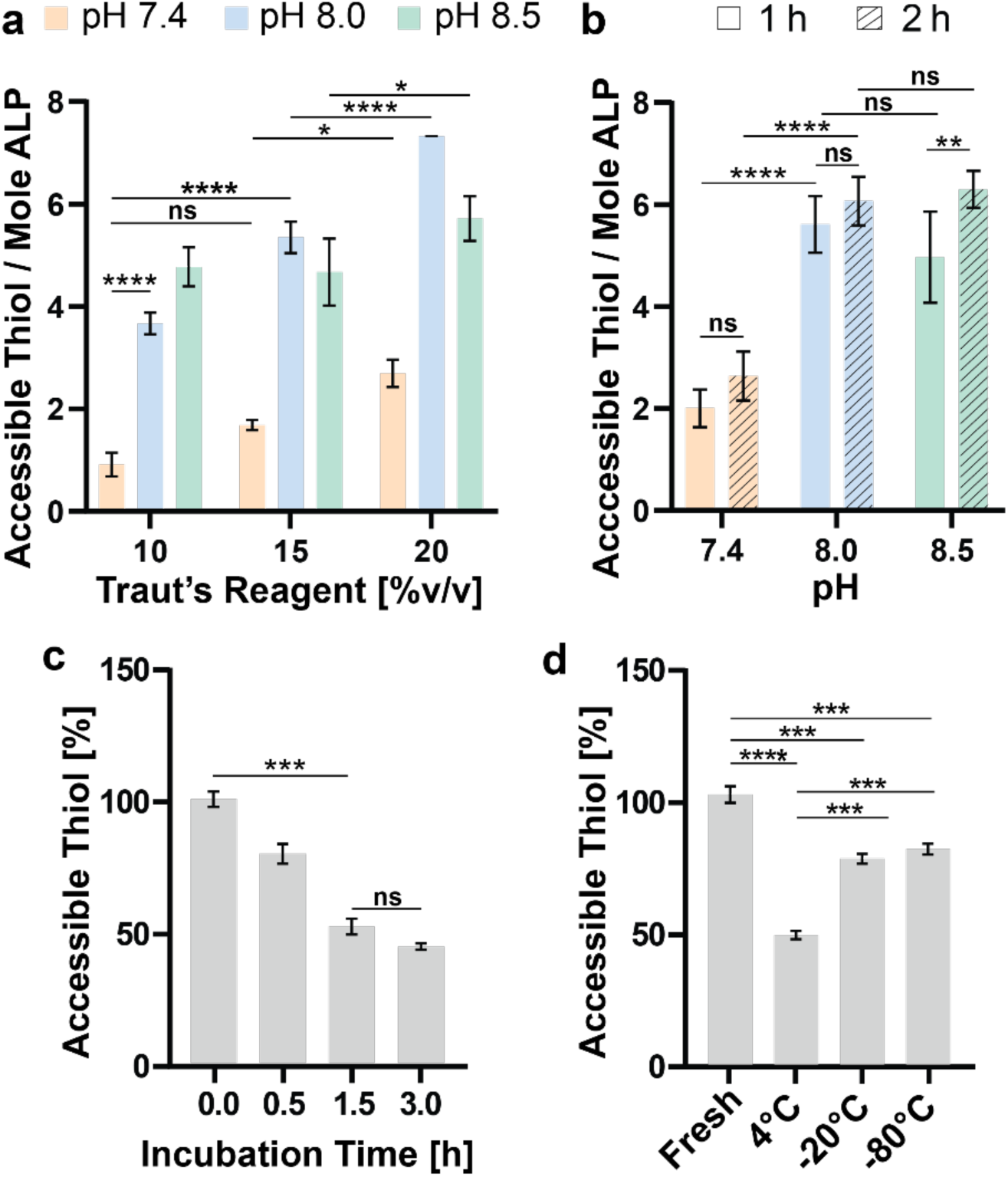
Efficiency of alkaline phosphatase (ALP) thiolation and stability of introduced free thiols. (a) Effect of Traut’s reagent concentration on the number of accessible thiols introduced per ALP at different reaction pH values at 20 °C. (b) Number of accessible thiols introduced per ALP enzymes as a function of reaction pH after 1 and 2 h incubation with Traut’s reagent at 20 °C. (c) Temporal stability of thiol groups introduced on ALP enzymes. (d) Relative stability of accessible thiols on ALP following storage at 4 °C, −20 °C, and −80 °C, normalized to freshly prepared samples. Data are presented as mean ± standard deviation (n = 3); statistical significance is indicated as *p < 0.05, **p < 0.01, ***p < 0.001, ****p < 0.0001 and ns: not-significant.

The degree of thiolation increased with Traut’s reagent concentration, but the extent of this increase depended strongly on reaction pH (Figure 2a). These experiments were carried out at pH 7.4, 8.0, and 8.5 to compare near-physiological conditions with mildly basic conditions expected to enhance amine reactivity^[31,32]^ and thereby a higher thiolation degree. At pH 7.4, thiolation remained limited across all conditions, yielding 1.1 ± 0.4, 2.0 ± 0.4, and 2.9 ± 0.5 thiol groups per ALP molecule at 10%, 15%, and 20% v/v Traut’s reagent, respectively. At pH 8.0, the effect of reagent concentration was more pronounced, with 10%, 15%, and 20% v/v yielding 3.8 ± 0.4, 5.6 ± 0.6, and 7.5 ± 0.2 thiols per ALP, respectively. At pH 8.5, 10%, 15%, and 20% v/v Traut’s reagent yielded 5.0 ± 0.6, 5.0 ± 0.9, and 5.9 ± 0.2 thiols per enzyme, respectively. Thus, although mildly basic conditions improved thiolation relative to pH 7.4, increasing the pH from 8.0 to 8.5 did not further enhance the measurable free thiol content. This observation likely reflects competing oxidation of newly introduced thiols at higher pH, where thiolate anions are more readily oxidized to disulfides despite ongoing lysine modification.^[33–35]^

We next evaluated the effect of reaction time on thiolation using 15% v/v Traut’s reagent at pH 7.4, 8.0, and 8.5 (Figure 2b). Extending the reaction time from 1 to 2 h increased thiolation at pH 7.4 by ∼30%, reaching 2.6 ± 0.5 thiol groups per ALP molecule. At pH 8.0, thiolation increased by ∼9% to 6.1 ± 0.5 thiols per enzyme, while at pH 8.5 a ∼26% increase was observed, reaching 6.3 ± 0.4 thiol groups per ALP molecule. Although the relative increase at pH 8.5 was more pronounced, extending the reaction time beyond 1 h provided only limited overall gains under mildly basic conditions, whereas thiolation at pH 7.4 remained markedly less efficient.

After functionalizing ALP with thiol groups, we performed a free thiol stability assay to determine the period over which the introduced thiols remain accessible at room temperature (Figure 2c). Detectable thiols dropped from 100% (immediately post-thiolation) to ∼75% at 0.5 h (post-purification), and to ∼45% by 3.0 h. This stability decay is consistent with the reported rapid oxidation of free thiols^[36–38]^ and highlights a narrow operational window between thiolation and downstream processes. Storage temperature also strongly influenced the stability of thiol groups introduced on ALP (Figure 2d). At 4 °C, only ∼50% of free thiols remained accessible relative to freshly prepared controls, whereas storage at −20 °C and −80 °C preserved ∼75–80% accessible free thiol groups.

Based on these results, standard thiolation conditions for subsequent experiments were set to pH 8.0, 20% v/v Traut’s reagent for 1 h at room temperature, yielding ∼6–8 thiols per ALP molecule (**Table S1**). Thiolated ALP was then stored at −80 °C in phosphate buffered saline (PBS) prior to use to limit oxidation and maintain batch-to-batch reproducibility.

### 2.2. ALP-functionalized PEG microgels are enzymatically active and assembly of injectable granular matrices

After optimization of thiolation conditions, we next incorporated SH-ALP into PEG-based microgels. Prior to microgel fabrication, we first verified that the PEG-Nb formulation reliably formed a stable hydrogel network under the photopolymerization conditions used by means of in situ photo-rheology on bulk formulations (**Figure 3a**). Before UV exposure, both storage (G′) and loss (G″) moduli were below the rheometer detection limit, consistent with properties of a liquid precursor. Upon UV exposure, G′ increased rapidly and plateaued within ∼60 s, reaching a final storage modulus of 2036 ± 4 Pa and tan(delta) of 0.002, confirming efficient crosslinking and network formation with mechanical properties consistent with previous reports.^[39,40]^

**Figure 3.**
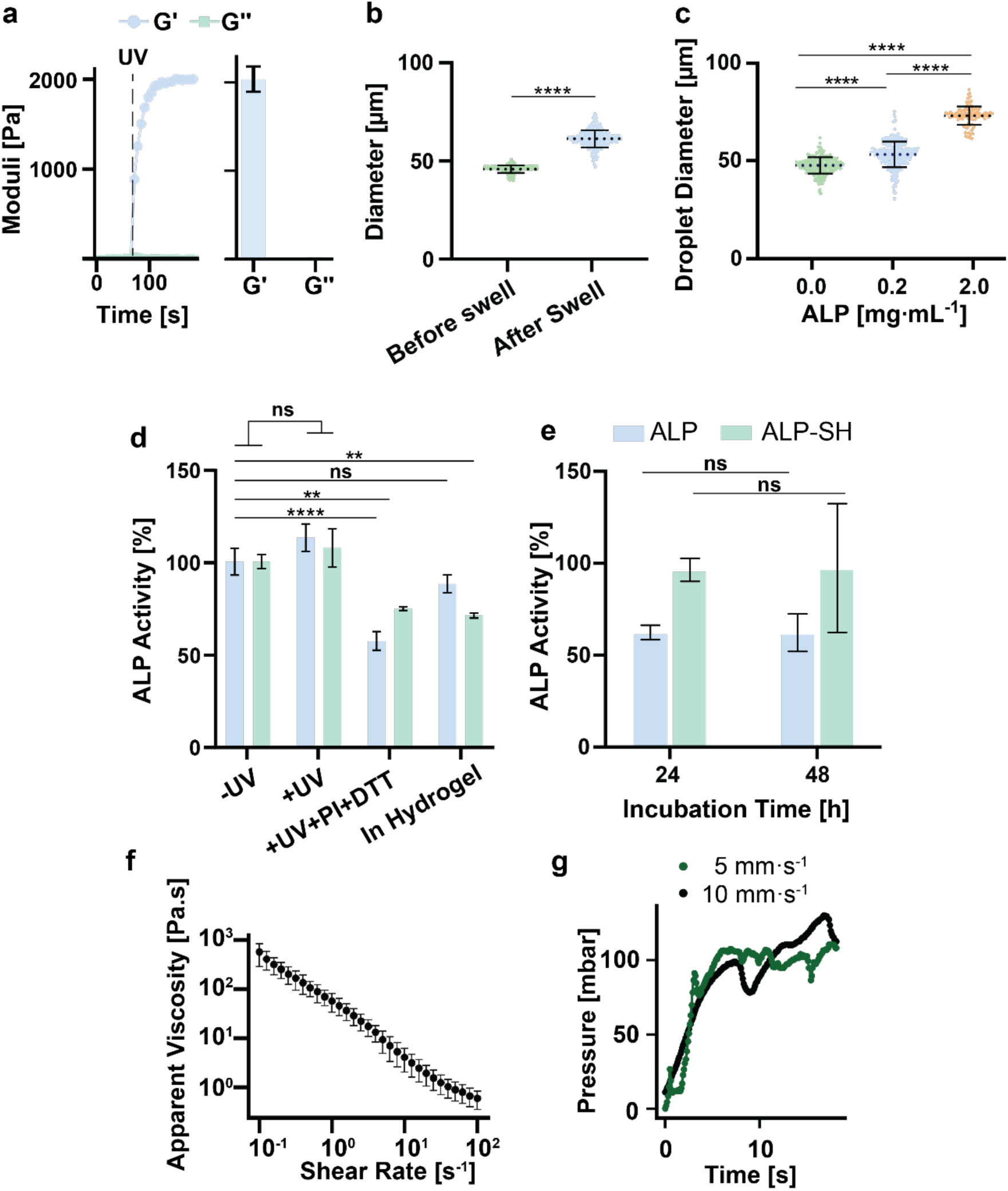
Physicochemical characterization of ALP-functionalized PEG microgels. (a) In situ photo-rheology of the PEG–norbornene (PEG-Nb) formulation showing the evolution of storage (G′) and loss (G″) moduli upon UV exposure (dashed line), with bar plots indicating average plateau moduli. (b) Microgel size before and after droplet demulsification in aqueous buffer, demonstrating hydrogel swelling. (c) Microdroplet diameter distributions as a function of alkaline phosphatase (ALP) concentration in the precursor solution measured in oil phase. (d) Relative activity of non-thiolated ALP (ALP) and thiolated ALP (SH-ALP) without UV exposure (−UV), with UV exposure (+UV), with UV exposure in the presence of lithium phenyl-2,4,6-trimethylbenzoylphosphinate (LAP) photoinitiator (+UV+PI+DTT), and after inclusion within PEG-Nb hydrogels. (e) Long-term activity retention of covalently immobilized SH-ALP compared to physically integrated ALP after incubation at 37 °C for 24 and 48 h. (f) Apparent viscosity of the microgel suspension as a function of shear rate measured by rotational rheology. (g) Representative pressure profiles obtained during capillary rheology measurements simulating injection of the microgel suspension through a nozzle with dimensions comparable to the microfluidic channels. Data are presented as mean ± standard deviation (n = 3 for panel a, d, e, f); statistical significance is indicated as *p < 0.05, **p < 0.01, ***p < 0.001, ****p < 0.0001 and ns: not significant.

We then proceeded with microgel formation by means of water-in-oil droplet microfluidics and characterized microgel size before and after droplet demulsification in aqueous buffers to evaluate their swelling behavior (Figure 3b). Microdroplets measured in oil exhibited a mean diameter of 42.9 ± 5.3 μm. Upon immersion of crosslinked microgels in aqueous medium, microgels displayed swelling, reaching an equilibrium diameter of 61.0 ± 5.5 μm within 4 hours.

We then examined the effect of ALP content in hydrogel precursor solution on droplet formation and resulting microgel size (Figure 3c). Increasing ALP concentration in the precursor solution from 0 to 0.2 mg·mL⁻¹ resulted in a modest increase in droplet diameter from 47.0 ± 4.1 μm to 52.5 ± 6.4 μm, while 2 mg·mL⁻¹ ALP yielded larger droplets of 72.0 ± 4.6 μm in diameter. This trend is consistent with previous reports regarding protein-induced changes in aqueous phase interfacial properties, which influence droplet breakup at the flow-focusing junction.^[41]^ Importantly, droplet monodispersity was maintained across all condition as the coefficient of variation for their size remained below 12%, indicating robust and reproducible microgel formation independent of enzyme loading within the tested concentration range.

Given the potential sensitivity of enzymes to UV irradiation and radical-mediated crosslinking, we next assessed the effect of the photochemical conditions used during microgel formation on ALP activity, defined as percent activity relative to unmodified ALP (Figure 3d). Following UV exposure alone (+UV), no significant change in activity was observed for both ALP and SH-ALP. In contrast, combined UV exposure, photoinitiator and crosslinker addition (+UV+PI+DTT) resulted in a pronounced activity reduction, particularly for ALP which retained 57.0 ± 5.0% activity, while SH-ALP retained 74.5 ± 1.0% activity. Notably, upon incorporation into the hydrogels, ALP exhibited 88.0 ± 4.9% activity which was similar to the non-irradiated control, while SH-ALP retained lower activity than ALP (70.7 ± 1.3%). Together, these findings indicate that although the applied UV exposure did not measurably compromise ALP activity in this system, radical-generating conditions can reduce enzyme function. Nonetheless, covalent conjugation of SH-ALP to the PEG network emerged as a highly effective strategy to preserve enzyme activity during microgel formation, with preserved activity remaining near non-irradiated controls.

We next evaluated whether covalently immobilized SH-ALP enzymes maintain their catalytic efficacy over the extended incubation periods required for on-chip mineralization. To this end, bulk hydrogels containing covalently anchored SH-ALP were compared with hydrogels in which non-thiolated ALP was physically entrapped and incubated at 37 °C for 24 and 48 h. (Figure 3e). hydrogels functionalized with SH-ALP maintained 95.3 ± 6.4% and 96.4 ± 35.1% of their initial activity after 24 and 48 h, respectively. In contrast, hydrogels containing non-thiolated ALP retained 61.1 ± 3.8% and 61.0 ± 10.3% activity after 24 and 48 h, respectively, indicating a consistent reduction in apparent activity that plateaued beyond 24 h. This sustained ALP activity loss in physically entrapped hydrogels likely reflects enzyme leaching from the matrix, deactivation, or a combination of both, whereas covalent conjugation approach preserved catalytic activity throughout the incubation period. Overall, these results support covalent tethering as the superior immobilization strategy for sustained catalysis under conditions relevant to on-chip mineralization.

To determine whether enzyme loading translated into functional mineralization output, microgels prepared with varying ALP concentrations (0, 0.2, and 2 mg·mL⁻¹) were incubated with 100 mM Ca-GP for 48 h and stained with Alizarin Red S to visualize calcium phosphate deposits (**Figure S1**). While ALP-free microgels showed no detectable mineral staining, ALP-conjugated microgels exhibited concentration-dependent increase in mineral deposition, with the strongest signal observed at 2 mg·mL⁻¹ ALP. These results confirm that the mineralization capacity of the microgels is governed by the degree of ALP conjugation and provides a functional basis for selecting the ALP concentration used in subsequent on-chip experiments.

Having established the activity and mineralization capacity of the ALP-functionalized microgels, we next assessed whether the microgel suspensions could be reliably injected through narrow channels for their on-chip integration as a granular matrix. This step was particularly critical given that granular materials often lead to nozzle clogging or phase separation during injection.^[42,43]^ To this end, we first characterized the flow behavior of the suspensions by rotational rheology, which revealed shear-thinning over a shear-rate range of 0.1–100 s⁻¹ (Figure 3f).

Such behavior is characteristic of jammed microgel suspensions and is commonly attributed to yielding and shear-induced particle rearrangement.^[42,44]^ Because rotational rheology alone does not fully capture injectability in hydrogel-based systems,^[43,45]^ we further evaluated extrusion behavior by capillary rheology using nozzles representative of the target microchannel dimensions in envisioned on-chip models (Figure 3g). The suspensions were extruded reproducibly at 104.3 ± 14.5 mbar and 102.3 ± 4.6 mbar for two extrusion speeds of 5 and 10 mms^-1^, respectively. These low extrusion pressure values and lack of adverse phenomena such as clogging or phase separation confirm suitability of this microgel suspension for injection into microfluidic chips. Pressure oscillations observed at the extrusion plateau likely reflect intermittent local rearrangements during confined granular flow, a feature characteristic of jammed suspensions.^[42,43]^

### 2.3. On-chip integration of ALP-functionalized microgels enables spatiotemporal control of mineral formation

To translate the enzymatically active microgels into a perfusable mineralization platform, we integrated these microgels into a microfluidic device to support assembly of a confined granular matrix. To this end, we designed a microfluidic chip comprising a central chamber (1 × 4 × 0.35 mm, width × length × height) to facilitate the assembly of a 3D matrix based on physical jamming of microgels. This central chamber was flanked by perfusion side channels separated by pillar arrays with 30 μm interpillar gaps (**Figure 4a**, **Figure S2**), providing two key functions: (i) physical retention of microgels within the central chamber and (ii) continuous lateral fluid exchange across the granular matrix. Finally, this platform features one inlet to the central channel allowing injection of microgel suspension and two access points per side channel for substrate delivery.

**Figure 4.**
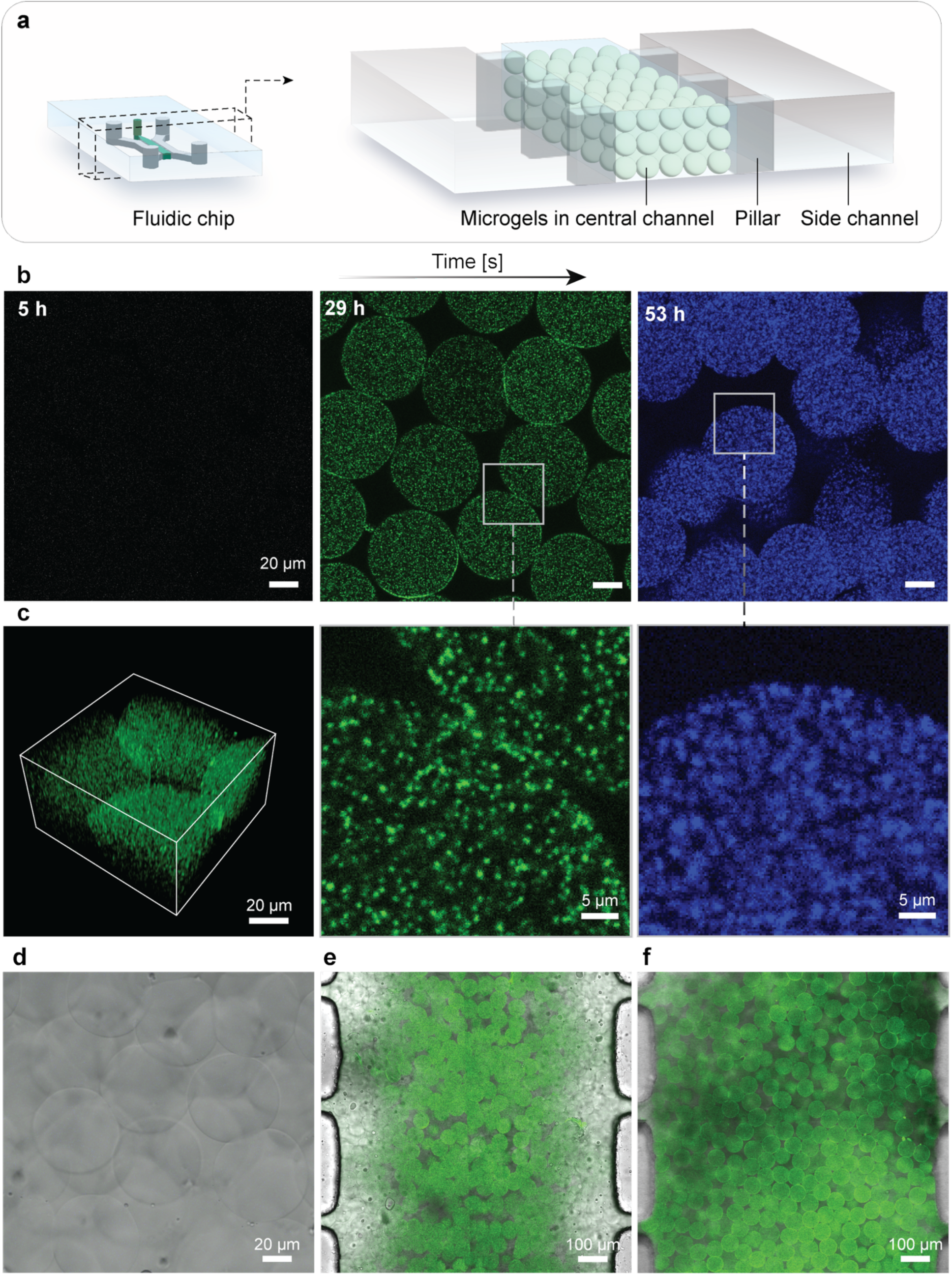
On-chip mineralization. (a) Schematic of the microfluidic chip illustrating the central channel packed with microgels, retained by a pillar array separating the central and side channels. (b) Confocal fluorescence images of the packed microgel bed during mineralization, showing the early-stage mineralization group after 5 h (Calcein Blue) and 29 h (Calcein Green), and the later-stage mineralization group after 53 h (Calcein Blue), with magnified views of the boxed regions at 29 h and 53 h (c) Three-dimensional confocal reconstruction of mineralized microgels within the packed bed. (d) Representative brightfield image of control group consisting of microgels without ALP. (e) Representative merged green and brightfield channels of fluorescent microscopy image captured for mineralization under semi-dynamic flow generated by rocker system. (f) Representative merged green and brightfield channels of fluorescent microscopy image captured for mineralization under static incubation conditions. Green fluorescence signal marks mineralized sections stained with Calcein Green.

Upon injection of microgel suspensions into the central chamber, microgels formed a 3D granular matrix composed of ∼20 vol% inter-microgel void space. This assembly was driven by pre-jamming the microgels before injection, followed by a second jamming step within the chip’s central channel, facilitated by the exit of the excess liquid phase into the side channels. To enable semi-dynamic substrate delivery using a rocker system, we developed 3D-printed custom reservoirs that were mounted onto the side-channel access points of the chip (**Figure S3**). The resulting assembly provides both adequate substrate volume and the geometry required for rocker-mediated perfusion of side channels, where the hydrostatic height difference between two reservoirs initiates flow. Subsequently, mineralization was initiated by perfusing Ca^2+^-GP through the side channels, thereby providing Ca²⁺ ions and a substrate for ALP-mediated hydrolysis to generate an inorganic phosphate source required for mineral formation. To monitor the temporal dynamics of mineral formation within the granular matrix, we employed sequential fluorochrome labeling, in which distinct fluorescent dyes are applied at successive time points to mark newly formed mineral deposits (Figure 4b). This approach, widely used to map bone mineral apposition rates in vivo,^[46]^ enabled us to distinguish early-nucleated mineral from mineral formed during subsequent phases. At 5 h, fluorescent signal was minimal, indicating limited early nucleation. By 29 h, distinct spot signals were visible throughout the granular matrix, indicating spatially distributed nucleation within the microgels rather than isolated precipitation on the microgels surface. At this stage, detected mineral features exhibited an average diameter of 1.0 ± 0.1 μm (n = 25,938). By 53 h, fluorescence intensity increased markedly across the construct, and the mean mineral feature diameter increased to 2.3 ± 0.2 μm (n = 14,680), indicating progressive growth and/or coalescence of mineral domains over time. The increase in feature size and intensity over time suggests sustained maturation of the mineral phase, rather than a short-lived nucleation burst. Three-dimensional confocal reconstruction (Figure 4c) confirmed volumetric mineral signal within individual microgels, demonstrating effective penetration of soluble precursors into the microgels’ polymeric network.

The observation of robust mineral deposition within PEG microgels is notable, given that PEG-based matrices have been reported to suppress calcium phosphate nucleation in the absence of nucleation-promoting sites.^[18]^ The observed mineralization suggests at least two possible scenarios: (i) nucleation occurs preferentially in the immediate vicinity of immobilized ALP, where localized phosphate generation creates microscale regions of elevated supersaturation, and/or (ii) sustained ALP activity increases phosphate levels more broadly within the microgel phase, such that nucleation can proceed throughout the matrix once supersaturation is reached. The dependence of mineral formation on enzyme activity was confirmed by analysis of ALP-free microgels under identical perfusion and mineralization conditions. In these controls, fluorochrome signal remained at or near background levels over matched time points (Figure 4d). This lack of measurable signals indicates that mineral formation in the experimental group is predominantly ALP-dependent, rather than spontaneous precipitation from media components.

To further probe how fluid flow influences mineralization dynamics, we compared rocker-mediated dynamic flow of substrate through side channels against static incubation. Microscopic observations of minerals formed under the dynamic flow condition revealed a spatial bias, with mineral formation appearing to occur preferentially near the central longitudinal axis of the granular matrix (Figure 4e). In contrast, the static condition produced a more uniform mineral distribution at matched early time points (Figure 4f). At later time points, mineralization expanded from the central axis toward pillars, indicating progressive chamber-wide growth once nucleation was established centrally.

We initially hypothesized that rocker-mediated flow would promote a more spatially homogeneous mineralization pattern than static conditions, as the resulting convective transport should overcome diffusional constraints and equilibrate substrate distribution. However, the observed spatial bias in mineralization location might suggest that convective disturbance of flow near pillars may disrupt the local accumulation of mineral precursors required for nucleation.^[47,48]^ In this context, the comparatively less disturbed flow along the central axis may permit earlier attainment of local supersaturation, resulting in preferential mineralization. These results highlight the potential role of local hydrodynamics in governing early mineral formation within perfused granular matrices. Consistent with this interpretation, confocal fluorescence imaging of the full channel length revealed pronounced longitudinal heterogeneity in mineral deposition, with certain regions exhibiting denser or earlier-onset deposits than others (**Figure S4**). This heterogeneity is likely due to local variations in substrate concentration gradients and interstitial flow velocity across the granular matrix, which further supports the transport-coupled nature of the mineralization process described above.

### 2.4. Mineral identity is consistent with a nanocrystalline apatite-like phase

To determine whether the on-chip mineralization supported by the granular matrix yielded a bone-relevant mineral phase, we examined the identity and crystallinity of the resulting mineral phase using complementary bulk and nanoscale analyses (**Figure 5**).

**Figure 5.**
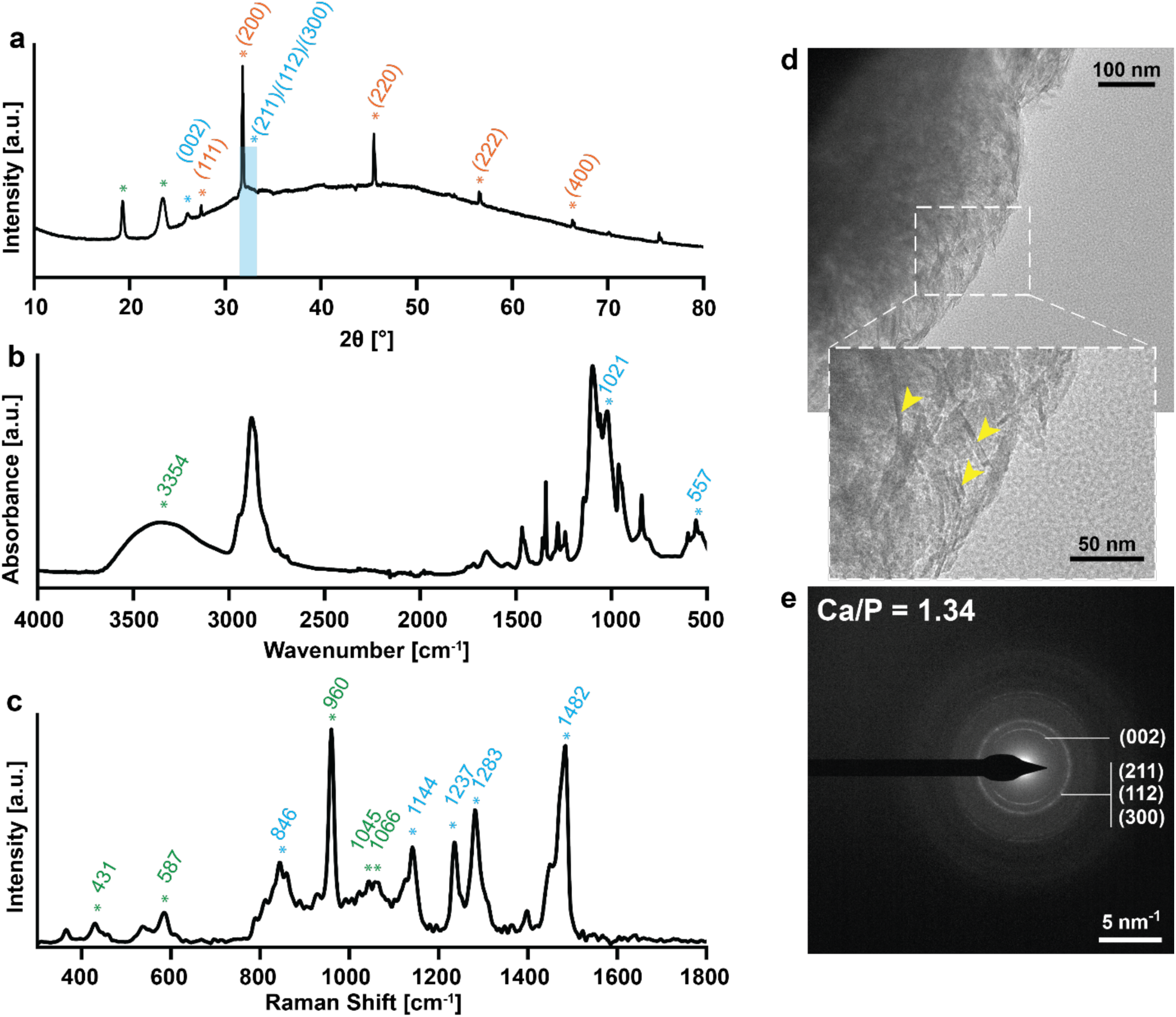
Physicochemical characterization of on-chip mineralized microgels. Color-coded annotations indicate mineral-related signals in blue, polymer-related signals in green, and residual halite-related XRD reflections in orange. (a) X-ray diffraction (XRD) pattern of mineralized microgels showing key reflections assigned to the mineral phase, polymeric phase, and residual halite. The blue shaded region marks the broad envelope assigned to overlapping (211)/(112)/(300) reflections of weakly crystalline hydroxyapatite. (b) Fourier-transform infrared (FTIR) spectrum of mineral deposits within the microgels. (c) Raman spectrum of mineralized microgels highlighting characteristic vibrational modes. (d) Transmission electron microscopy (TEM) image of mineral morphology together with a higher magnification view of the needle-shaped minerals indicated by yellow arrows. (e) Selected area electron diffraction (SAED) pattern of the mineral phase with the corresponding Ca/P ratio determined by inductively coupled plasma optical emission spectroscopy (ICP-OES). All measurements were performed on dried microgels after 3 days of mineralization at 37 °C.

X-ray diffraction (XRD; Figure 5a) was used to assess the crystalline structure of the mineral formed within the microgels. Evidence of mineralization was observed as a broad reflection near 2θ ≈ 26° and a broad envelope underlying the sharp peak at 2θ ≈ 31.9°, corresponding to the (002) and merged (211)/(112)/(300) reflections of weakly crystalline hydroxyapatite, respectively. The pronounced broadening of both the 26° reflection and the envelope underlying the 32° peak indicates small crystallite size and low crystallinity. The sharp peak at 2θ ≈ 31.9° itself is attributed to the (200) reflection of halite (NaCl), arising from crystallization of residual PBS during drying of the hydrogels; this assignment is confirmed by additional sharp NaCl reflections at 2θ ≈ 27.5° (111), 45.6° (220), 56.6° (222), and 66.3° (400). The sharp reflections at 2θ ≈ 19.3° and 23.5° are attributed to the (120) and (132) planes of crystalline PEG, which forms as the polymer chains fold and pack upon dehydration of the hydrogel.^[49]^ Despite the interference from the buffer salts, the characteristic broadening of the underlying mineral reflections is consistent with the formation of an early-stage, apatite-like calcium phosphate phase with limited crystallinity.^[50]^ Such phases, which often contain an amorphous precursor component, are commonly observed in enzyme-driven biomimetic and biological mineralization systems.^[51–55]^

Fourier-transform infrared spectroscopy (FTIR; Figure 5b) provided complementary information on the chemical composition of the mineral phase. Characteristic phosphate absorption bands were observed at 962 cm⁻¹ (ν₁ PO₄³⁻ symmetric stretching), as a resolved doublet at 599 and 557 cm⁻¹ (ν₄ PO₄³⁻ bending modes), and in the 1020–1100 cm⁻¹ region (ν₃ PO₄³⁻ asymmetric stretching), the latter partially overlapping with C–O–C vibrations of the polymer matrix.^[17,56,57]^ The resolution of the ν₄ PO₄³⁻ doublet indicates a degree of short-range crystalline order, consistent with a poorly crystalline but structurally defined apatite-like phase.^[55]^ A broad absorption between 3000 and 3500 cm⁻¹ reflects structural water and hydroxyl-containing groups,^[58]^ and weak bands in the 1415–1500 cm⁻¹ region are consistent with minor carbonate (ν₃ CO₃²⁻) incorporation into the apatite lattice, a hallmark of biologically relevant bone-like apatitic phases.^[59–61]^

Raman spectroscopy further supported the presence of a carbonated apatitic phase (Figure 5c). These Raman spectra showed a dominant ν₁ PO₄³⁻ peak near ∼960 cm⁻¹, the diagnostic marker of apatitic calcium phosphates, with additional phosphate-associated modes in the 400–600 cm⁻¹ (ν₂/ν₄) and 1000–1100 cm⁻¹ (ν₃) ranges. Furthermore, low-intensity carbonate-associated features in the 1200–1500 cm⁻¹ interval support partial carbonate substitution, a hallmark of biological bone mineral.^[62,63]^

At the nanoscale, transmission electron microscopy (TEM; Figure 5d) revealed irregular, electron-dense aggregates tens of nanometers in size, composed of fine needle-shaped sub-features. To assess the local crystallinity of the formed minerals at microscale/nanoscale, selected area electron diffraction (SAED; Figure 5e) was employed. The diffraction pattern yielded a continuous ring pattern rather than discrete spots, confirming a polycrystalline assembly of randomly oriented nanocrystallites. The measured d-spacings of 3.42 and ∼2.6 Å were assigned to the (002) and overlapping (211)/(112)/(300) reflections, respectively, with an additional outer ring at ∼1.96 Å indexed to the (222) reflection. The fact that the (211)/(112)/(300) reflections appear as a single unresolved envelope, together with the moderate arcing of the rings, is indicative of small crystallite size and limited long-range order, consistent with the XRD reflections.^[64]^ An additional diffuse inner ring at larger d-spacings (∼4–6 Å) is attributed to the surrounding organic matrix and is not associated with the mineral phase.

Elemental analysis yielded a Ca/P molar ratio of 1.34 (Figure 5e, inset), below the stoichiometric value of hydroxyapatite (1.67) and consistent with a calcium-deficient, non-stoichiometric apatite.^[57,65]^ Such non-stoichiometry is typical of early-stage biomimetic apatites formed under kinetically controlled conditions and suggests that the PEG microenvironment and perfusion-controlled substrate delivery promote a mineral composition closer to the biological counterpart than to a fully matured stoichiometric hydroxyapatite phase.^[66,67]^

Although a transient ACP precursor is frequently reported in enzyme-driven and matrix-mediated mineralization,^[54,55,57]^ our data indicate that the mineral phase is already predominantly nanocrystalline at the timepoint examined. This conclusion is evidenced by the resolved ν₄ PO₄³⁻ doublet in the FTIR spectrum and discrete SAED rings. Under sustained ionic supply and controlled transport, extended maturation is expected to drive further crystallite growth and transition toward more ordered apatitic phases,^[18,55,57,68]^ motivating future long-term studies to map phase-evolution kinetics on chip.

## 3. Conclusion and outlook

In this work, we engineered an enzymatically active granular matrix that enables on-chip modeling of near-physiological, bone-like mineralization. Our results confirm that PEG-based microgels covalently conjugated with ALP effectively act as catalytic microcompartments and, when assembled in a microfluidic chamber, form a perfusable granular matrix. By coupling compartmentalized ALP chemistry with perfusion, the system enables in situ calcium phosphate nucleation and deposition, producing a carbonated, calcium-deficient, apatite-like phase with a low crystallinity, consistent with an early-stage biomimetic mineral state.

Accordingly, this platform provides a mechanistically defined acellular framework for interrogating ALP-driven mineralization in a perfusable microfluidic environment. By enabling independent control over enzyme loading, substrate delivery, and local transport, the on-chip platform supports spatially localized, time-resolved mineralization under defined conditions, while excluding confounding effects associated with cell-state heterogeneity, matrix remodeling, and coupled biological signaling.

Importantly, the modularity of microgel-based matrix design employed in this work creates future opportunities to increase the biological complexity of on-chip systems. For instance, enzymatically active microgels could be combined with cell-laden compartments to generate hybrid biosynthetic models in which catalytic and cellular modules are tuned independently. Such systems may provide a useful route to dissect how localized enzymatic activity, transport, and cell-mediated regulation interact during (patho)physiological tissue mineralization. More broadly, this work identifies compartmentalized enzymatic function within perfusable granular matrices as a promising design principle for engineering mineralized tissue models with programmable composition and spatial organization.

## 4. Experimental section

### Thiolation of alkaline phosphatase

50 mg ALP (Phosphatase, Alkaline from bovine intestinal mucosa, Sigma Aldrich) was dissolved in 4 mL of 0.5 M 4-(2-hydroxyethyl)-1-piperazineethanesulfonic acid (HEPES) buffer (pH 8.0) to achieve a final concentration of 10 mg mL⁻¹. Traut’s reagent stock solution was prepared at 2 mg mL⁻¹ in deionized water immediately before use. To initiate thiolation, 1 mL of Traut’s reagent stock (20% v/v final concentration) was added to the ALP solution. The reaction mixture was protected from light with aluminum foil and incubated at room temperature (21°C) for 1 hour under gentle rotation (5 rpm). The thiolated ALP (SH-ALP) was purified by size exclusion using Zeba Spin Desalting Columns (7 kDa molecular weight cut-off, MWCO, Thermo Fisher Scientific, USA) pre-equilibrated with PBS (pH 7.4). The column was centrifuged at 1000 × g for 2 minutes to remove storage solution, washed three times with 5 mL PBS, and the SH-ALP sample was applied and eluted by centrifugation. Purified SH-ALP was aliquoted and stored at −80°C until use.

### Quantification of accessible thiol groups

Accessible thiol content was quantified using Ellman’s assay.^[69]^ Ellman’s reagent solution was prepared by dissolving 4 mg 5,5′-dithiobis(2-nitrobenzoic acid) (DTNB) in 1 mL dimethyl sulfoxide (DMSO). A cysteine standard curve was prepared with concentrations ranging from 0 to 1.5 mM in 0.5 M HEPES buffer (pH 8.0). In a 96-well plate, 2 μL of Ellman’s reagent solution was added to each well, followed by 100 μL of reaction buffer and 10 μL of standard or sample. The plate was incubated at room temperature for 15 minutes protected from light, and absorbance was measured at 412 nm using a BioTek Synergy HTX multimode plate reader (Agilent Technologies, USA). Thiol concentration was calculated from the standard curve, and thiolation degree was expressed as moles of thiol per mole of ALP.

### ALP activity assay

ALP activity was measured using para-nitrophenyl phosphate (pNPP) as substrate, following previous reports.^[11]^ Substrate solution (5 mM pNPP) was prepared in assay buffer (10 mM HEPES, 150 mM NaCl, pH 7.4). A 4-nitrophenol (4-NP) standard curve (0–250 μM) was prepared in assay buffer. In a 96-well plate, samples (2–3 μL) were added to wells containing assay buffer to reach 100 μL total volume, followed by 100 μL substrate solution. The plate was sealed with adhesive film and incubated at 37°C for 60 minutes. The reaction was stopped by adding 100 μL of 0.3 M NaOH, and absorbance was measured at 405 nm. ALP activity was calculated from the 4-NP standard curve and expressed as percent activity relative to unmodified ALP.

### Preparation of PEG-norbornene hydrogel precursor solution

A 5% (w/v) PEG-Nb hydrogel precursor solution was prepared by dissolving 4-arm PEG-Nb (Creative PEGWorks, North Carolina, USA) in PBS buffer. Dithiothreitol (DTT, Sigma Aldrich) stock solution (2.2 mg mL⁻¹ in buffer) and lithium phenyl-2,4,6-trimethylbenzoylphosphinate (LAP, Sigma Aldrich) stock solution (4 mg mL⁻¹ in buffer) were prepared fresh. Thawed SH-ALP was added to the PEG-Nb solution at a final concentration of 1 mg mL⁻¹, followed by DTT as the crosslinker and to reduce any disulfide bonds formed during storage, and LAP (final concentration 2.27 mM). The solution was mixed by vortexing, centrifuged briefly (1000 × g, 2 min) to remove bubbles, and protected from light until use. For visualization of microdroplets or microgels, fluorescein isothiocyanate-dextran (FITC-dextran, 1 mg mL⁻¹) was included into the precursor solution for microdroplet imaging, whereas sulfo-Cy5–tetrazine was conjugated to the PEG microgels post-crosslinking via incubation at 4 °C overnight.

### Fabrication of microgels

Microgels were fabricated using a flow-focusing droplet microfluidic platform (Figure S2a). To ensure consistent droplet formation, the polydimethylsiloxane (PDMS) channels were functionalized with a hydrophobic coating by flowing Rain-X (ITW Global Brands, USA) through the device at 10 µL·min^-1^ for 15 min, followed by thermal treatment at 65 °C for 45 min. The oil phase consisted of Novec™ HFE-7500 Engineered Fluid (3M™, USA) supplemented with 1 wt% 008-FluoroSurfactant in HFE7500 (RAN Biotechnologies, USA). Both phases were filtered through 0.2 µm units before loading into syringes.

The solutions were infused into the chip using a syringe pump (Chemyx 5000, USA). After stabilizing the flow at 10 µL·min^-1^, the flow rates were adjusted to 5 µL·min^-1^ for the aqueous phase and 15 µL·min^-1^ for the oil phase to generate monodisperse droplets. The collected emulsion was UV-cured (365 nm, 15 mW·cm⁻²) for 2 min to induce crosslinking. To recover the microgels, the oil phase was removed, and the emulsion was broken by adding 20% perfluoro-1-octanol (PFO) in a 2:1 (v/v) ratio with the sample. The resulting microgels were washed with PBS, centrifuged, and stored at 4 °C until use.

### Microfluidic chip fabrication

PDMS microfluidic chips used for droplet generation (Figure S2a), and mineralization (Figure S2b) were fabricated by soft lithography. Master molds were designed using Autodesk Fusion 360 and 3D printed using a DLP printer (Sonic Mini 8K s, Phrozen, Taiwan) with Aqua Hyperfine resin (Phrozen, Taiwan). Following fabrication, the molds were washed with isopropanol (IPA), air-dried, and post-cured in a UV chamber (Formlabs Form Cure V1, Formlabs, Somerville, MA, USA) for 180 min at room temperature. To ensure structural stability and remove residual monomers, the 3D-printed molds underwent a secondary thermal treatment at 120 °C for 8 h. Once cooled, the mold surfaces were functionalized using trichloro(1H,1H,2H,2H-perfluorooctyl)silane (Sigma-Aldrich) for 45 min to facilitate PDMS release.

PDMS (Sylgard 184, Dow Corning) was prepared by mixing the elastomer base and curing agent in a 10:1 (w/w) ratio. The mixture was degassed in a vacuum desiccator until all air bubbles were removed. The prepolymer was then cast onto the 3D-printed molds and thermally crosslinked at 65 °C for 2 h. Post-curing, the PDMS replicas were peeled from the molds, and fluidic access ports were created using a biopsy punch. Finally, the PDMS layers and glass coverslips were treated with air plasma (Harrick, USA) and immediately brought into contact to form a permanent covalent bond.

### Rheological characterization

#### Rotational rheology

Viscoelastic properties of bulk hydrogels and microgel suspensions were characterized using oscillatory rheometry (AR2000 Advanced Rheometer, TA Instruments, USA). To study gelation kinetics, hydrogel precursor solutions were loaded between a parallel plate geometry (8 mm diameter, 0.5 mm gap) and a custom-made in situ UV exposure module. Time sweeps were performed at 0.1% strain and 1 Hz frequency to monitor gelation kinetics during which UV exposure (15 mW·cm⁻²) photocrosslinked the precursor in situ. Flow sweeps were conducted at shear rate of 0.1–100 s⁻¹ at 0.1% strain.

#### Capillary rheology

Extrusion/injection performance of microgel suspension was assessed using X-TRUDE,^[43]^ a custom-made capillary rheology system. After mounting microgel-loaded syringes onto the syringe holders of X-TRUDE system, the plunger was positioned in contact with syringe pistons. Extrusion tests were conducted with an applied linear extrusion rate of 10 and 5 mm·s^-1^ using a 6 mm long (20G, inner diameter 0.6 mm) cylindrical nozzle. Each extrusion step was maintained at these constant shear rates until a steady-state pressure plateau was reached. Real-time pressure values were used for further analysis. All measurements were conducted at 21 °C.

#### In situ assembly of microgels

To prepare the granular hydrogel compositions, the fabricated microgels were first pre-jammed via centrifugation at 10,000 relative centrifugal force (RCF) for 5 min. The supernatant was carefully aspirated using an air displacement pipette, yielding a volume fraction of approximately 75%. The pre-jammed microgel suspension was then loaded into a 25 µL positive displacement pipette and injected into the central channel of the mineralization chip. Injection was maintained until the microgels reached the channel terminus, where the confinement of the channel geometry further compressed the microgels into a uniform, jammed assembly with a final volume fraction of approximately 80–85%.

#### On-chip mineralization

For the mineralization process, reservoirs were fitted to all channel inlets and outlets of the mineralization chip (Figure S3), and each reservoir was filled with 150 µL of 10 mM calcium glycerophosphate (Ca-GP) solution in deionized water and solution pH was adjusted to 8.0. To maintain a humidified environment and prevent matrix dehydration, the devices were placed in 140 mm Petri dishes alongside a secondary vessel containing 10 mL of PBS. The assemblies were sealed with Parafilm and incubated at 37 °C to facilitate in situ mineralization. Every 24 h, the Ca-GP solution was fully aspirated from all inlet and outlet reservoirs and replaced with fresh 150 µL of 10 mM Ca-GP solution.

Dynamic and static mineralization conditions were investigated. For static mineralization, the sealed assemblies were placed in an incubator at 37 °C for 24 and 48 h. For dynamic mineralization, the sealed assemblies were placed on a rocker system inside the incubator at 37 °C, operating at a tilt angle of 15° and a frequency of 1 RPM, for 24 and 48 h.

#### Incorporation of fluorochromes into microgels for tracking mineral deposition kinetics

Matrix-integrated mineralization chips were initially incubated with 400 µL of 10 mM Ca-GP solutions per chip at pH 8.0 to initiate mineralization. Chips were divided into two groups to visualize mineral deposition at early and later stages.

For early-stage mineralization, chips were first incubated with Ca-GP for 5 h. The channels were then washed twice with deionized (DI) water. Calcein Blue solution (4 mg/mL, 25 µL per reservoir) was introduced and incubated for 15 min to label mineral deposits formed during the initial 5 h mineralization period. Unbound dye was removed by four 5-min washes with DI water. The chips were then re-incubated with 10 mM Ca-GP for an additional 24 h. Calcein Green solution (4 mg/mL, 25 µL per reservoir) was subsequently introduced and incubated for 15 min to label minerals deposited during the second mineralization period.

For later-stage mineralization, chips were incubated with Ca-GP for 53 h from the start of the experiment. The channels were then washed twice with DI water. Calcein Blue solution (4 mg/mL, 25 µL per reservoir) was introduced and incubated for 15 min to label mineral deposits formed over the 53 h mineralization period. Unbound dye was removed by four 5-min washes with DI water.

#### Fluorescence microscopy and image analysis

Microgel imaging was performed using a laser scanning confocal microscope (LSM 880, Zeiss, Germany). For microgel size analysis, non-mineralized microgels were imaged using a 10× objective with a 633 nm excitation laser and a 650–750 nm emission filter to detect Cy5 fluorescence. Z-stacks were acquired at 5 µm intervals and reconstructed using Fiji (open-source, ImageJ distribution).^[70]^ Microgel segmentation was performed using Cellpose (v3.0.5, cyto2 cp3 model),^[71]^ and the resulting masks were imported into Fiji for particle analysis. Diameters were calculated from measured areas assuming circular geometry.

For mineral deposition analysis, mineralized microgels stained with Calcein Green (excitation 488 nm, emission 500–550 nm) and Calcein Blue (excitation 405 nm, emission 420–480 nm) were imaged using a 20× objective. Z-stacks were acquired at 5 µm intervals and reconstructed in Fiji (for detailed analysis see Supplementary Methods).

#### Physicochemical characterization of mineralization

Fourier Transform Infrared Spectroscopy (FTIR) spectra were recorded on a spectrometer (Spectrum Two™, PerkinElmer, USA) equipped with an attenuated total reflectance (ATR) diamond crystal accessory. Mineralized microgels were dried at room temperature for 5 days prior to analysis. Spectra were acquired from 400 to 4000 cm⁻¹ with a resolution of 2 cm⁻¹ and 32 scans per spectrum. Background spectra were collected before each sample measurement. Spectra were baseline corrected and normalized using PerkinElmer Spectrum software.

Raman spectra were recorded using a WITec alpha 300R confocal Raman microscope (Oxford Instruments, UK) with a 532 nm excitation laser operated at 40 mW, and a Zeiss EC Epiplan-Neofluar DIC 50×/0.8 NA objective lens. Spectra were acquired over the range of 267– 2392 cm⁻¹ with an acquisition time of 6 s and 20 accumulations per spectrum. One spectrum was collected per sample. The raw Raman spectra were baseline-corrected using the WITec 5 Project software. Powder X-ray diffraction (XRD) measurements were performed using a Panalytical Empyrean diffractometer (Malvern Panalytical, UK) operating in Bragg-Brentano geometry using Cu Kα radiation (45 kV, 40 mA) and equipped with a PIXcel3D 1x1 detector. To this end, samples were dispersed as a thin layer on a zero-background (557) silicon wafer. A continuous scan was made over the 5° < 2θ < 90° range with a step size of 0.013°.

Transmission electron microscopy (TEM, Talos F200C G2, Thermo Scientific, USA) operated at 200 kV, combined with selected area electron diffraction (SAED), was used to characterize the morphology and crystal structure of formed minerals. TEM imaging was performed at a magnification of 92,000×. SAED patterns were recorded at a camera length of 850 mm. Dried microgels were dispersed onto a carbon-coated copper TEM grid before the measurements. Inductively coupled plasma–optical emission spectrometry (ICP-OES; Spectro Ametek, Germany) was used to quantify elemental concentrations. Detection wavelengths were 422.673 nm for calcium (Ca) and 177.495 nm for phosphorus (P). Calibration was performed using a five-point standard series: STD 1 (Ca: 5.0002 ppm, P: 10.002 ppm), STD 2 (Ca: 0.9903 ppm, P: 1.9939 ppm), STD 3 (Ca: 0.5182 ppm, P: 0.3837 ppm), with quality control samples (QC = 1.000 ppm) measured at Ca: 0.9830–0.9931 ppm and P: 0.9987–0.9998 ppm, confirming calibration accuracy. Dried mineralized microgels harvested from 24 chips were dissolved in 10% (v/v) nitric acid (Sigma-Aldrich) and incubated for 30 minutes at 37 °C to ensure complete digestion. Following dissolution, the solution was diluted with demineralized water to a final nitric acid concentration of 2% (v/v). Measurements were performed in triplicate (n = 3). Based on the measured Ca and P concentrations, the Ca/P molar ratio was calculated.

### Statistical Analysis

All experiments were performed in triplicate (n = 3) unless otherwise stated. Results are reported as mean ± standard deviation. Statistical significance was assessed by one-way analysis of variance (ANOVA) with Tukey’s post-hoc test for multiple comparisons or Student’s t-test for pairwise comparisons using GraphPad Prism 10 software (GraphPad Software, San Diego, CA, USA) and custom programming scripts using Python 3.12 (Python Software Foundation, open-source). Differences were considered statistically significant at p < 0.05. Significance levels are indicated as: *p < 0.05, **p < 0.01, ***p < 0.001, ****p < 0.0001 and ns: non-significant.

## Acknowledgements

This work was funded by Hypatia Grant (BoneChipPredict) from Radboud University Medical Center and VIDI Grant from The Dutch Research Council (NWO, grant no. 22190). M.D. acknowledges a Marie Skłodowska-Curie Actions (MSCA) Individual Global Fellowship (grant no. 891016) from European Commission. The authors thank Prof. David J. Mooney for valuable discussions and scientific input during the early stages of this work. The authors acknowledge Dr. Anqi Chen for early guidance in microfluidic device design and fabrication.

## Data availability statement

The data that support the findings of this study are available from the corresponding author upon reasonable request.

