## Supplementary Information for "Engineering an Enzymatically Active Granular Matrix for On-Chip Modeling of Bone-Like Mineralization"

1. Supplementary methods

##### **Image-based quantification of fluorescent puncta in microgels**

### *Adaptive thresholding with watershed segmentation for time point 29h:* Mineral particle segmentation was performed using a custom Python pipeline (Python 3.12; OpenCV 4.11.0; scikit-image 0.26.0; SciPy 1.17.0). Each greyscale image was first enhanced using Contrast Limited Adaptive Histogram Equalisation (CLAHE; clip limit = 3.0, tile grid = 8 × 8 pixels) to normalise local contrast variations across scaffold cross-sections. A light Gaussian blur (σ ≈ 1.5 pixels, kernel = 3 × 3) was applied to suppress single-pixel noise while preserving mineral boundaries.

Bright mineral deposits were identified by Gaussian-weighted adaptive thresholding (block size = 51 pixels, offset constant *C* = −8), which classifies each pixel as foreground or background relative to the intensity-weighted mean of its local neighborhood. This approach accommodates the gradual intensity variation across individual scaffold filaments and between filament cross-sections. The resulting binary mask was cleaned by morphological opening with an elliptical structuring element (3 × 3 pixels) to eliminate isolated noise pixels.

To separate touching or partially overlapping mineral particles, a marker-controlled watershed algorithm was applied. The Euclidean distance transform of the binary foreground mask was computed, and local maxima (minimum inter-peak distance = 5 pixels) within the distance map were identified as seed markers. These markers were dilated by one pixel and labelled as initial regions, then the watershed transform was applied to the inverted distance map to delineate individual particle boundaries. Segmented objects with a projected area below 4 pixels² (sub-resolution noise) or above 500 pixels² (merged clusters or imaging artefacts) were excluded from further analysis. Equivalent diameter was converted to micrometers using a calibration factor for the 29 h dataset, 3.77 pixels = 1 µm, reflecting acquisition with the 20× objective.

Summary statistics — including mean, median, standard deviation, interquartile range, and full range — were computed for each layer or Z-position. Size distributions were visualized as probability density histograms, box plots, and cumulative distribution functions.

*Top-hat and Otsu-based puncta analysis with size-filtered ROI quantification for time point 53h:* Fluorescent puncta were quantified from 2D microscopy images using a custom pipeline implemented in Python (Python 3.12; OpenCV 4.11.0; scikit-image 0.26.0; SciPy 1.17.0). For color images, the green channel was used as the analysis channel; for grayscale images, the native intensity image was used. All analyses were performed on cropped fields at identical processing settings within each dataset.

To enhance punctate signal and suppress slowly varying background, images were smoothed with a Gaussian filter (σ = 1 pixel), followed by white top-hat filtering with a disk structuring element (radius = 6 pixels). A global Otsu threshold was then applied to obtain a binary puncta mask. Binary masks were refined by removing small, connected components and filling small holes (minimum area threshold = 6 pixels).

Connected puncta were segmented into individual regions from the cleaned binary mask using connected-component labeling, followed by size-based post-filtering to remove residual debris and oversized regions likely representing merged objects. Objects smaller than 6 pixels were excluded in all analyses. The maximum allowed area was set per image set to account for magnification-dependent object scale (A_max_ = 120). Segmentation quality was verified visually by superimposing detected region boundaries on the original analysis-channel image.

For each detected object, area (pixels²), equivalent diameter (pixels), and centroid coordinates were extracted. Equivalent diameter was converted to micrometers using a calibration factor for 53 h dataset, 8.08 pixels = 1 µm, reflecting acquisition with the 10× objective. Because puncta-size distributions were right-skewed/non-Gaussian, results were summarized using the median and interquartile range (IQR; 25th–75th percentile). For pooled analysis of the three Image 15 fields, per-object equivalent diameters from all three images were combined prior to summary statistic calculation.

To evaluate sensitivity to size filtering, the minimum object area threshold was varied (4, 6, 8, 10, and 12 pixels) while all other parameters were held constant; 6 pixels was used as the primary threshold for reporting.

2. Supplementary figures

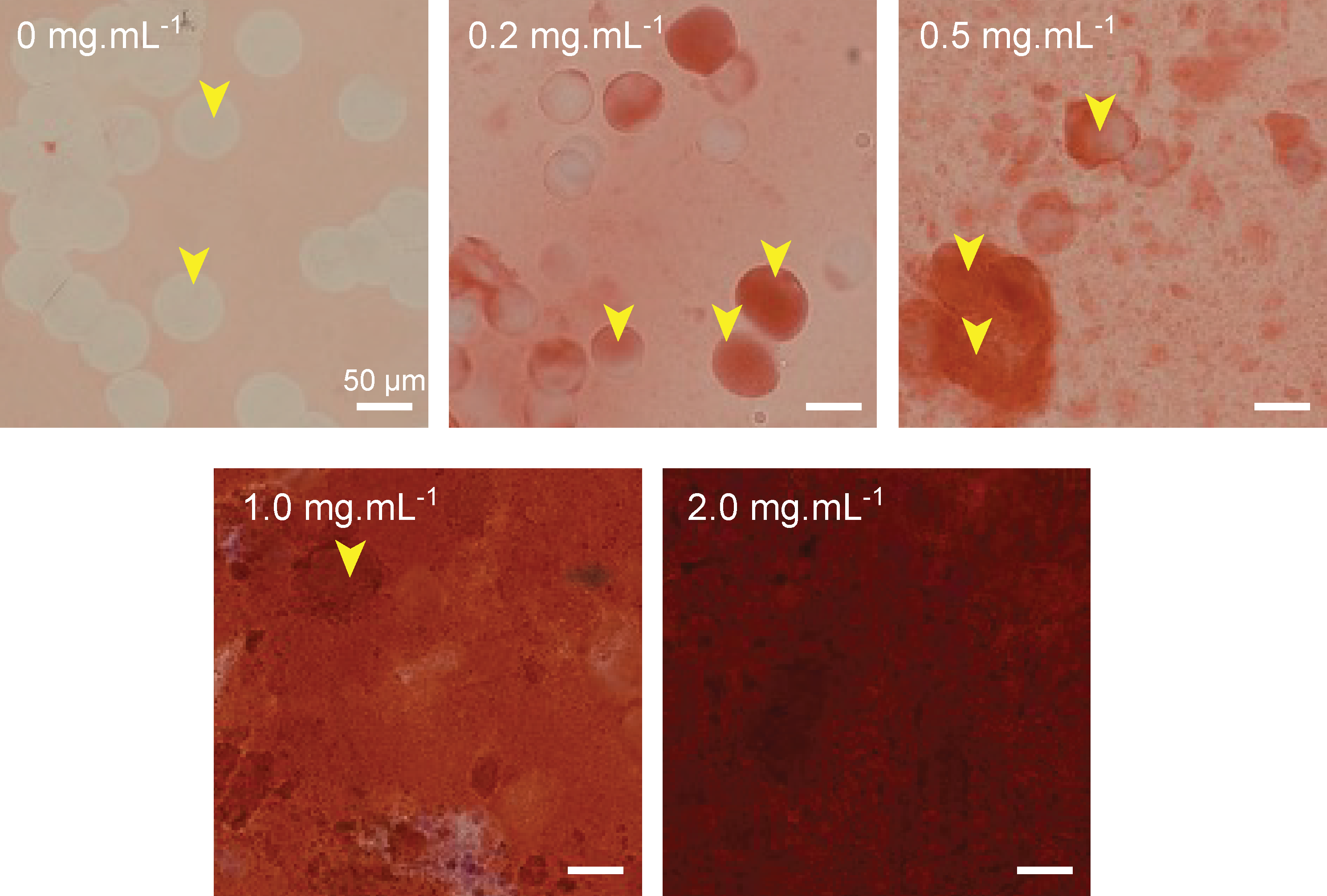

Figure S1. Representative images of microgels fabricated with varying ALP concentrations in the hydrogel precursor solution (0, 0.2, and 2 mg/mL) after incubation with 100 mM calcium glycerophosphate for 48 hours, stained with Alizarin Red S to visualize calcium phosphate deposits. Arrows indicate microgels.

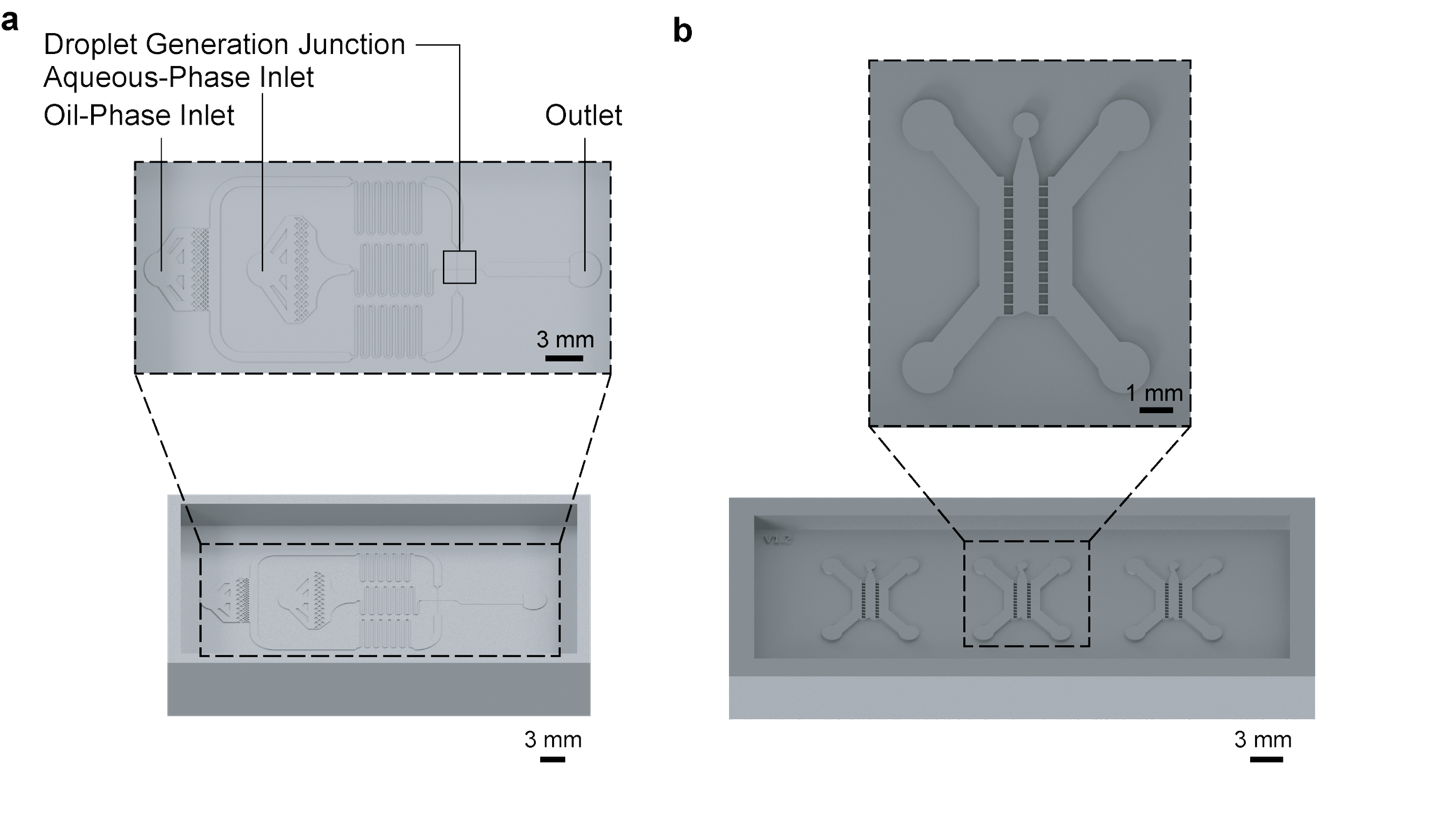

Figure S2. (a) 3D computer aided design (CAD) rendering of droplet generation platform fabrication mold with a close-up top view. (b) 3D CAD rendering of mineralization platform fabrication mold containing three chips with a close-up top view of central chip.

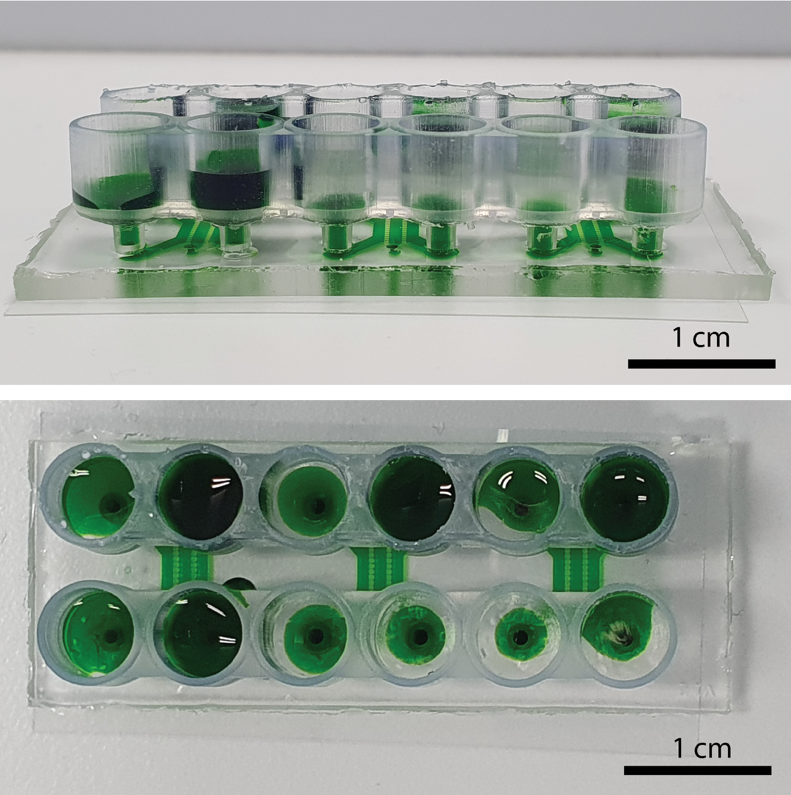

Figure S3. Custom-designed snap-on reservoirs attached to the PDMS mineralization chip inlets and outlets, filled with green food dye solution for demonstration purposes.

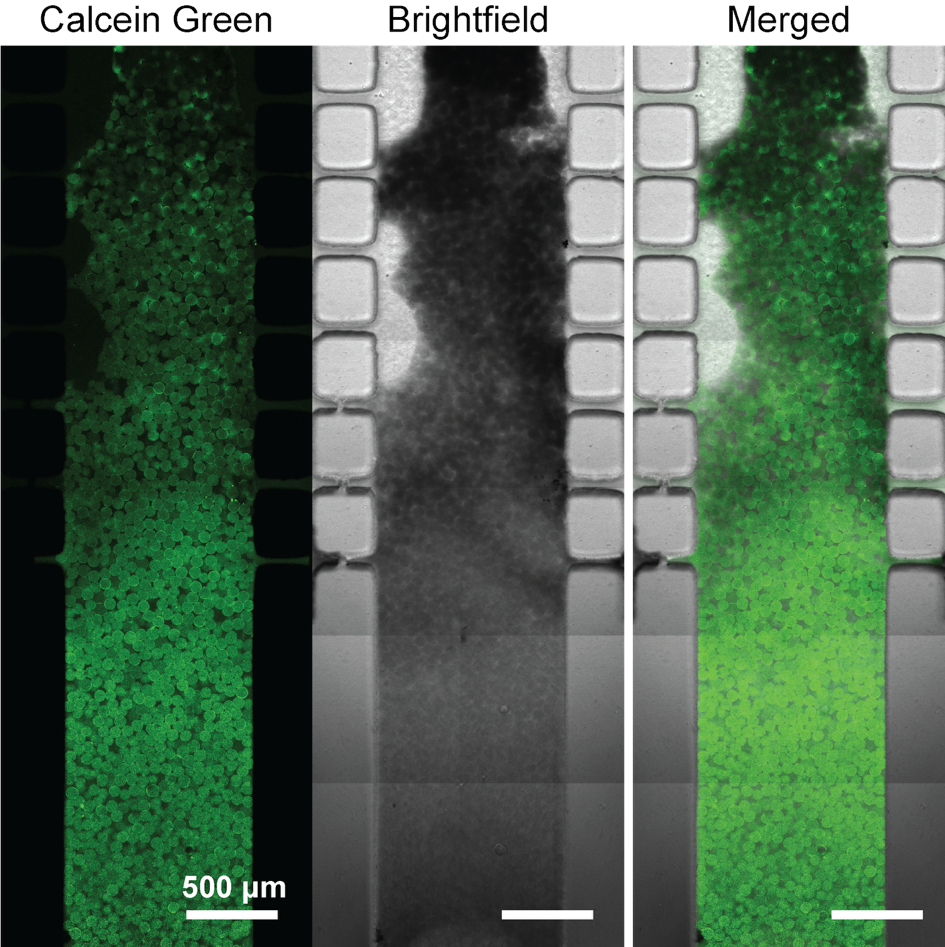

Figure S4. Non-uniform distribution of minerals throughout the central channel. Confocal fluorescence images showing spatial heterogeneity of mineral deposition across the granular matrix, revealing preferential mineralization patterns influenced by local transport conditions. Green color indicates formed minerals.

3. Supplementary tables

Table S1. Summary of thiolation optimization experiments

| **Parameter** | **Range Tested** | **Optimal Value** | **Notes** |
| --- | --- | --- | --- |
| pH | 7.4, 8.0, 8.5 | 8.0 | Balance of efficiency and thiol stability |
| Traut's reagent | 10, 15, 20 [% v/v] | 20 [% v/v] | Higher = more thiols |
| Reaction time | 1, 2 h | 1 h | Longer time minimal gain |
| Temperature | 21, 37 [°C] | 21 [°C] | Higher T = more oxidation |
| Thiols/ALP molar ratio | 2–8 | 6–8 (at pH 8) | Depends on conditions |
| Activity retained relative to ALP | 70–85 [%] | ~80 [%] | After thiolation |
